# GSA-Genie: a web application for gene set analysis

**DOI:** 10.1101/125443

**Authors:** Zhe Zhang, Deanne Taylor

**Affiliations:** Department of Biomedical and Health Informatics The Children’s Hospital of Philadelphia

## Abstract

Gene set analysis is often used to interpret results from upstream analysis through predefined gene sets that are linked to biological features such as cell cycle or tumorgenesis. Gene sets have been defined in the literature via various criteria and are archived by numerous databases. We compiled over 2.3 million gene sets from 17 sources, and made them accessible through a web application, GSA-Genie. Selected gene sets can be analyzed online using one of 16 statistical methods. These methods can be grouped into two strategies: test of gene set over-representation within a gene list, or comparison of a gene-level statistics between gene set and background. GSA-Genie operates on a Shiny web server, hosted in a cloud instance within Amazon Web Services. GSA-Genie offers a broad selection of gene sets and statistical methods comparing to existing tools. GSA-Genie is freely available at http://gsagenie.awsomics.org.

## 1 Introduction

Gene sets are groups of genes with one or more shared biological features, such as those involved in the same signaling pathways or activated together by a genetic perturbation. For example, the Gene Ontology defines gene sets based on collective knowledge about genes [1]; KEGG maps genes to canonical metabolic pathways [2], and iProClass classifies genes according to protein sequence, structure, and function [3]. Furthermore, integrative databases like MsigDb [4] collect gene sets from primary databases and stores them in a consistent format.

Gene set analysis (GSA) is often used by researchers to link their own results to a predefined gene set, and hence the biological features associated to that gene set. There are two common strategies of running GSA. The first tests if a gene set is over-represented in a project-specific gene list provided by the analyst. The gene list could be selected based on differential gene expression, burden test of mutation frequency, or any other analysis. DAVID is a popular online GSA tool using this stragegy [5]. The second strategy requires a gene-level statistic, such as the fold change of differential expression or the p value of burden test. The statistics is used for a two-group comparison of means or distributions between each gene set and the background. GSEA [6] is a standalone program that applies this strategy. A variety of statistical tests have been developed to analyze different types of gene-level statistics (t-like, p-like, etc.), and The R/Bioconductor package *piano* [7] implemented 11 such methods.

GSA-Genie is a web application and an alternative to the tools and databases mentioned above. It provides a one-stop, comprehesive solution of running GSA by allowing users to choose from 16 statistical methods and over 2.3 million gene sets sourced from 17 original databases.

## 2 Functionality

GSA-Genie sets up an analysis in 3 steps: select gene sets; upload user inputs; and choose method (Figure 1A).

**Figure 1.**
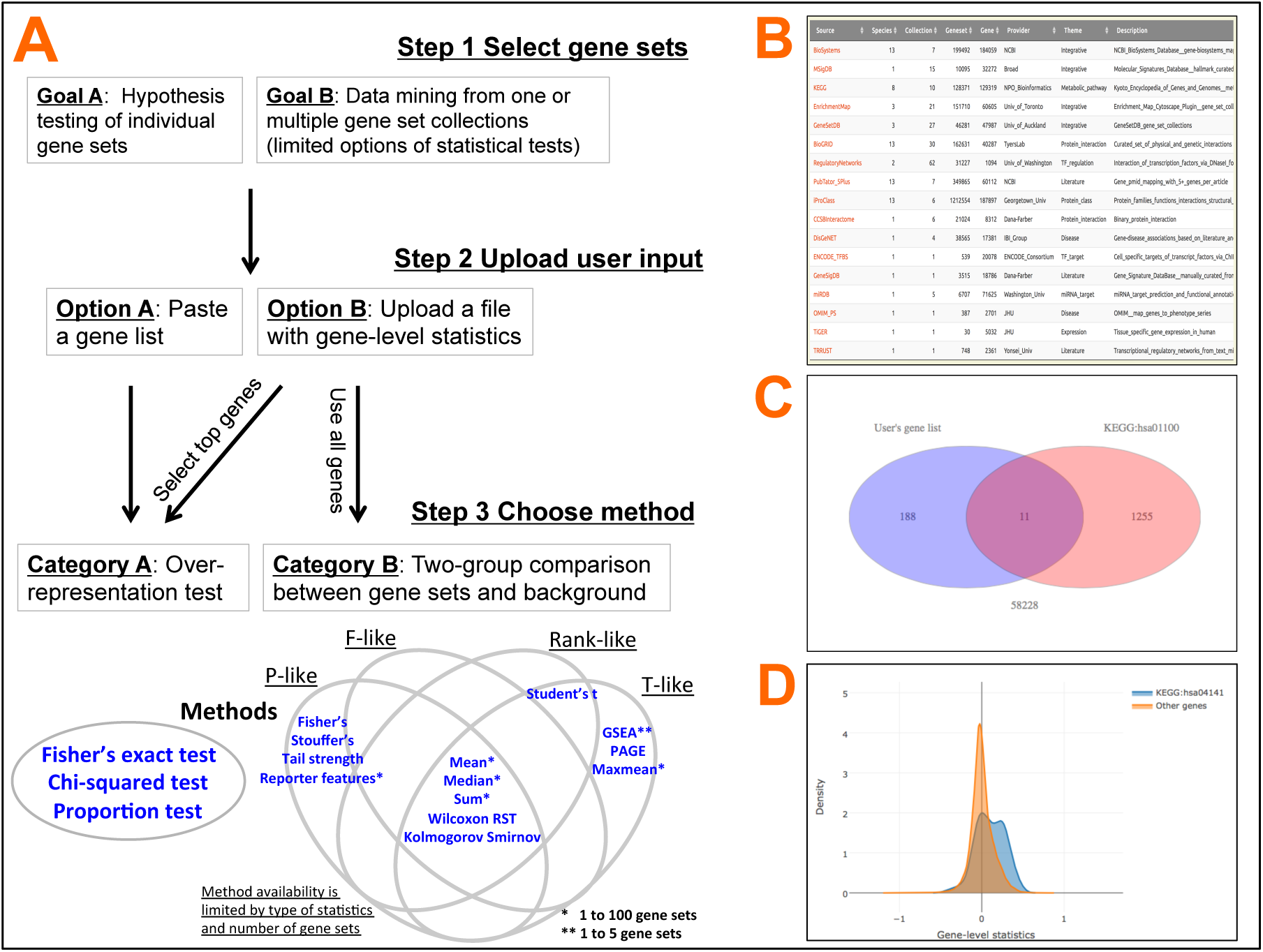
GSA-Genie. A). The overall workflow of using GSA-Genie. B). The summary table of gene set sources. C). Over-representation test of gene set analysis. D). Two-group comparison of gene set analysis.

### 2.1 Step 1: select gene sets

Gene sets were collected from 17 sources (Figure 1B): BioSystems [8], MSigDB, KEGG, Enrichment Map [9], GeneSetDB [10], TF regulatory networks [11], PubTator [12], iProClass, Interactome [13], DisGeNET [14], BioGRID [15], ENCODE [16], GeneSigDB [17], miRDB [18], OMIM [19], TiGER [20], and TRRUST [21]. All gene sets are consistently annotated and structured into collections that can be browsed by specifying species-source-collection, such as mouse-KEGG-pathway. Three types of gene identifiers are available for all gene sets: NCBI ID, Official sybmol, and Ensembl ID. Gene sets can be downloaded for offline analysis in several formats including the .gmt format used by GSEA. Instead of using whole collections of gene sets, users can perform hypothesis testing on one or a few selected gene sets of interest to save runtime and reduce multiple-testing burden.

### 2.2 Step 2: upload input

GSA-Genie accepts two types of user input: an ID-only gene list or a table of N rows of genes and at least one column of gene-level statistics. Both must use one of the three recognizable gene identifiers. When the input is a gene list, the GSA must be over-representation test. Users also have the option to upload the testing background to which the gene list will be compared. When gene-level statistics is uploaded, users can use the statistics to select top genes for overrepresentation test or use all genes in the table for two-group comparison between each gene set and all other genes in the uploaded table.

### 2.3 Step 3: choose method

Three methods are available for over-representation tests: Fisher’s exact (hypergeometric), Chi-squared, and proportion tests. Users can choose all known genes or all genes in selected gene set collections as background if they did not upload their own.

The R/Bioconductor *piano* package implements 11 statistical methods for two-group comparison of gene-level statistics, such as PAGE [22] and tail strength [23]. GSA-Genie added two more methods, Student’s t and Kolmogorov-Smirnov tests, for mean and distribution comparisons respectively. These methods offers analytical flexibility corresponding to different types of gene-level statistics. For example, Fisher’s combined test is for p-like statistics only while GSEA requires t-like statistics. Non-parametric methods, such as Wilcoxon rank sum test, are more robust and applicable to all types of statistics. GSA-Genie also provides options to transform one type of statistics to another. For example, re-scaling plus adding directionality information can convert a p-like statistics to a t-like one. GSA-Genie also implements a configurable methods-vs-dataset size gatekeeping to avoid testing a large number of gene sets using slower methods. For example, GSEA, which is much slower than the other methods, is only available for simultaneously testing 5 or less gene sets. This limit can be adjusted when running GSA-Genie on a local instance.

## 3 Results

GSA-Genie presents the results online immediately after an analysis is completed. No matter which method was used, each tested gene set will have a p value and corresponding false discovery rate. For the over-representation test, odds ratio and enrichment percentage are also included in the results.

Overlapping between a selected gene set and user’s gene list can be plotted as a venn diagram (Figure 1C). For two-group comparison, the results often include a method-specific test statistics, such as the t statistics of Studen’s t test and the enrichment score of GSEA. Methods using the directionality information of t-like statistics also report extra directional p values. Users can also select a gene set to visualize its density distribution versus the background distribution (Figure 1D).

## References

1: Ashburner M, Ball CA, Blake JA, Botstein D, Butler H, Cherry JM, Davis AP, Dolinski K, Dwight SS, Eppig JT, Harris MA, Hill DP, Issel-Tarver L, Kasarskis A, Lewis S, Matese JC, Richardson JE, Ringwald M, Rubin GM, Sherlock G. Gene ontology: tool for the unification of biology. The Gene Ontology Consortium. Nat Genet. 2000 May;25(1):25–9. PubMed PMID: 10802651; PubMed Central PMCID: PMC3037419.

2: Ogata H, Goto S, Sato K, Fujibuchi W, Bono H, Kanehisa M. KEGG: Kyoto Encyclopedia of Genes and Genomes. Nucleic Acids Res. 1999 Jan 1;27(1):29–34. PubMed PMID: 9847135; PubMed Central PMCID: PMC148090.

3: Wu CH, Xiao C, Hou Z, Huang H, Barker WC. iProClass: an integrated, comprehensive and annotated protein classification database. Nucleic Acids Res. 2001 Jan 1;29(1):52–4. PubMed PMID: 11125047; PubMed Central PMCID: PMC29833.

4: Liberzon A, Subramanian A, Pinchback R, Thorvaldsdóttir H, Tamayo P, Mesirov JP. Molecular signatures database (MSigDB) 3.0. Bioinformatics. 2011 Jun 15;27(12):1739–40. doi:10.1093/bioinformatics/btr260. Epub 2011 May 5. PubMed PMID: 21546393; PubMed Central PMCID: PMC3106198.

5: Huang da W, Sherman BT, Lempicki RA. Systematic and integrative analysis of large gene lists using DAVID bioinformatics resources. Nat Protoc. 2009;4(1):44–57. doi: 10.1038/nprot.2008.211. PubMed PMID: 19131956.

6: Subramanian A, Tamayo P, Mootha VK, Mukherjee S, Ebert BL, Gillette MA, Paulovich A, Pomeroy SL, Golub TR, Lander ES, Mesirov JP. Gene set enrichment analysis: a knowledge-based approach for interpreting genome-wide expression profiles. Proc Natl Acad Sci U S A. 2005 Oct 25;102(43):15545–50. Epub 2005 Sep 30. PubMed PMID: 16199517; PubMed Central PMCID: PMC1239896.

7: Väremo L, Nielsen J, Nookaew I. Enriching the gene set analysis of genome-wide data by incorporating directionality of gene expression and combining statistical hypotheses and methods. Nucleic Acids Res. 2013 Apr;41(8):4378–91. doi: 10.1093/nar/gkt111. Epub 2013 Feb 26. PubMed PMID: 23444143; PubMed Central PMCID: PMC3632109.

8: Geer LY, Marchler-Bauer A, Geer RC, Han L, He J, He S, Liu C, Shi W, Bryant SH. The NCBI BioSystems database. Nucleic Acids Res. 2010 Jan;38(Database issue):D492–6. doi: 10.1093/nar/gkp858. Epub 2009 Oct 23. PubMed PMID: 19854944; PubMed Central PMCID: PMC2808896.

9: Merico D, Isserlin R, Stueker O, Emili A, Bader GD. Enrichment map: a network-based method for gene-set enrichment visualization and interpretation. PLoS One. 2010 Nov 15;5(11):e13984. doi: 10.1371/journal.pone.0013984. PubMed PMID: 21085593; PubMed Central PMCID: PMC2981572.

10: Araki H, Knapp C, Tsai P, Print C. GeneSetDB: A comprehensive meta-database, statistical and visualisation framework for gene set analysis. FEBS Open Bio. 2012 Apr 17;2:76-82. doi: 10.1016/j.fob.2012.04.003. Print 2012. PubMed PMID: 23650583; PubMed Central PMCID: PMC3642118.

11: Neph S, Stergachis AB, Reynolds A, Sandstrom R, Borenstein E, Stamatoyannopoulos JA. Circuitry and dynamics of human transcription factor regulatory networks. Cell. 2012 Sep 14;150(6):1274–86. doi: 10.1016/j.cell.2012.04.040. Epub 2012 Sep 5. PubMed PMID: 22959076; PubMed Central PMCID: PMC3679407.

12: Wei CH, Kao HY, Lu Z. PubTator: a web-based text mining tool for assisting biocuration. Nucleic Acids Res. 2013 Jul;41(Web Server issue):W518–22. doi: 10.1093/nar/gkt441. Epub 2013 May 22. PubMed PMID: 23703206; PubMed Central PMCID: PMC3692066.

13: Venkatesan K, Rual JF, Vazquez A, Stelzl U, Lemmens I, Hirozane-Kishikawa T, Hao T, Zenkner M, Xin X, Goh KI, Yildirim MA, Simonis N, Heinzmann K, Gebreab F, Sahalie JM, Cevik S, Simon C, de Smet AS, Dann E, Smolyar A, Vinayagam A, Yu H, Szeto D, Borick H, Dricot A, Klitgord N, Murray RR, Lin C, Lalowski M, Timm J, Rau K, Boone C, Braun P, Cusick ME, Roth FP, Hill DE, Tavernier J, Wanker EE, Barabási AL, Vidal M. An empirical framework for binary interactome mapping. Nat Methods. 2009 Jan;6(1):83–90. doi: 10.1038/nmeth.1280. Epub 2008 Dec 7. PubMed PMID: 19060904; PubMed Central PMCID: PMC2872561.

14: Piñero J, Bravo À, Queralt-Rosinach N, Gutiérrez-Sacristán A, Deu-Pons J, Centeno E, García-García J, Sanz F, Furlong LI. DisGeNET: a comprehensive platform integrating information on human disease-associated genes and variants. Nucleic Acids Res. 2017 Jan 4;45(D1):D833–D839. doi: 10.1093/nar/gkw943. Epub 2016 Oct 19. PubMed PMID: 27924018; PubMed Central PMCID: PMC5210640.

15: Chatr-Aryamontri A, Oughtred R, Boucher L, Rust J, Chang C, Kolas NK, O’Donnell L, Oster S,Theesfeld C, Sellam A, Stark C, Breitkreutz BJ, Dolinski K, Tyers M. The BioGRID interaction database:2017 update. Nucleic Acids Res. 2017 Jan 4;45(D1):D369–D379. doi: 10.1093/nar/gkw1102. Epub 2016 Dec 14. PubMed PMID: 27980099; PubMed Central PMCID: PMC5210573.

16: Gerstein MB, Kundaje A, Hariharan M, Landt SG, Yan KK, Cheng C, Mu XJ, Khurana E, Rozowsky J, Alexander R, Min R, Alves P, Abyzov A, Addleman N, Bhardwaj N, Boyle AP, Cayting P, Charos A, Chen DZ, Cheng Y, Clarke D, Eastman C, Euskirchen G, Frietze S, Fu Y, Gertz J, Grubert F, Harmanci A, Jain P, Kasowski M, Lacroute P, Leng J, Lian J, Monahan H, O’Geen H, Ouyang Z, Partridge EC, Patacsil D, Pauli F, Raha D, Ramirez L, Reddy TE, Reed B, Shi M, Slifer T, Wang J, Wu L, Yang X, Yip KY, Zilberman-Schapira G, Batzoglou S, Sidow A, Farnham PJ, Myers RM, Weissman SM, Snyder M. Architecture of the human regulatory network derived from ENCODE data. Nature. 2012 Sep 6;489(7414):91–100. doi: 10.1038/nature11245. PubMed PMID: 22955619; PubMed Central PMCID: PMC4154057.

17: Culhane AC, Schröder MS, Sultana R, Picard SC, Martinelli EN, Kelly C, Haibe-Kains B, Kapushesky M, St Pierre AA, Flahive W, Picard KC, Gusenleitner D, Papenhausen G, O’Connor N, Correll M, Quackenbush J. GeneSigDB: a manually curated database and resource for analysis of gene expression signatures. Nucleic Acids Res. 2012 Jan;40(Database issue):D1060–6. doi: 10.1093/nar/gkr901. Epub 2011 Nov 21. PubMed PMID: 22110038; PubMed Central PMCID: PMC3245038.

18: Wong N, Wang X. miRDB: an online resource for microRNA target prediction and functional annotations. Nucleic Acids Res. 2015 Jan;43(Database issue):D146–52. doi: 10.1093/nar/gku1104. Epub 2014 Nov 5. PubMed PMID: 25378301; PubMed Central PMCID: PMC4383922.

19: Amberger JS, Bocchini CA, Schiettecatte F, Scott AF, Hamosh A. OMIM.org: Online Mendelian Inheritance in Man (OMIM^®^), an online catalog of human genes and genetic disorders. Nucleic Acids Res. 2015 Jan;43(Database issue):D789–98. doi: 10.1093/nar/gku1205. Epub 2014 Nov 26. PubMed PMID: 25428349; PubMed Central PMCID: PMC4383985.

20: Liu X, Yu X, Zack DJ, Zhu H, Qian J. TiGER: a database for tissue-specific gene expression and regulation. BMC Bioinformatics. 2008 Jun 9;9:271. doi: 10.1186/1471-2105-9-271. PubMed PMID:18541026; PubMed Central PMCID: PMC2438328.

21: Han H, Shim H, Shin D, Shim JE, Ko Y, Shin J, Kim H, Cho A, Kim E, Lee T, Kim H, Kim K, Yang S, Bae D, Yun A, Kim S, Kim CY, Cho HJ, Kang B, Shin S, Lee I. TRRUST: a reference database of human transcriptional regulatory interactions. Sci Rep. 2015 Jun 12;5:11432. doi: 10.1038/srep11432. PubMed PMID: 26066708; PubMed Central PMCID: PMC4464350.

22: Kim SY, Volsky DJ. PAGE: parametric analysis of gene set enrichment. BMC Bioinformatics. 2005 Jun 8;6:144. PubMed PMID: 15941488; PubMed Central PMCID: PMC1183189.

23: Taylor J, Tibshirani R. A tail strength measure for assessing the overall univariate significance in a dataset. Biostatistics. 2006 Apr;7(2):167–81. Epub 2005 Dec 6. PubMed PMID: 16332926.

